# Active Gαi/o mutants accelerate breast tumor metastasis via the c-Src pathway

**DOI:** 10.1101/2023.01.16.524334

**Authors:** Cancan Lyu, Aarzoo K Bhimani, William T Draus, Ronald Weigel, Songhai Chen

**Author notes:** Corresponding author: Songhai Chen, MD, PhD, The Department of Neuroscience and Pharmacology, Roy J. and Lucille A. Carver College of Medicine, University of Iowa, Iowa City, Iowa. 2-250 Bowen Science Building, 51 Newton Road, Iowa City, IA 52242.

## Abstract

Constitutively active mutations in the Gα_i2_ and Gα_oA_ subunits of heterotrimeric G proteins have been identified in several human cancers including breast cancer, but their functional significance in tumorigenesis and metastasis has not been well characterized. In this study, we show that expression of the constitutively active Gα_oA_R243H and Gα_i2_R179C mutants alone was insufficient to induce mammary tumor formation in mice. However, in transgenic mouse models of breast cancer induced by Neu expression or PTEN loss, we found that the Gα_i2_R179C mutant enhanced spontaneous lung metastasis while having no effect on primary tumor initiation and growth. Additionally, we observed that ectopic expression of the Gα_oA_R243H and Gα_i2_R179C mutants in tumor cells promote cell migration *in vitro* as well as dissemination into multiple organs *in vivo* by activating c-Src signaling. Thus, our study uncovers a critical function of Gα_i/o_ signaling in accelerating breast cancer metastasis via the c-Src pathway. This work is clinically significant, as it can potentially pave the way to personalized therapies for patients who present with active Gα_i/o_ mutations or elevated Gα_i/o_ signaling by targeting c-Src to inhibit breast cancer metastasis.

## Introduction

Breast cancer is the most commonly diagnosed cancer in women (1). As of 2020, breast cancer surpassed lung cancer to become the most prevalent form of cancer worldwide (1). In the United States, approximately 1 in 8 women is expected to be diagnosed with breast cancer in her lifetime (1). Despite major and rapid advances in therapeutic approaches, breast cancer remains the leading cause of death from malignant tumors among women. Metastasis is the major cause of tumor mortality. While the average 5-year survival rate for localized breast cancer is 99%, it falls to only 29% for advanced breast cancer diagnoses with distant metastasis (www.cancer.org). Since currently available therapies have very limited efficacy in treating metastatic disease, further research into the mechanisms of tumor metastasis is crucial. Understanding these mechanisms on a molecular level is critical for developing novel therapies that will ultimately improve the survival rates of advanced breast cancer patients.

G protein coupled receptors (GPCRs) are the largest cell surface receptor family that regulates diverse cellular processes and functions (2). GPCRs transmit signals through heterotrimeric G proteins that consist of Gα, Gβ and Gγ subunits. Based on sequence and functional similarity of Gα subunits, heterotrimeric G proteins are divided into four major subclasses: G_i/o_, G_s_, G_q/11_ and G_12/13_ (3, 4). GPCRs regulate diverse cellular processes including cell recognition, proliferation and motility. Not surprisingly, dysregulated GPCR and G protein signaling has been implicated in cancer initiation, progression, and metastasis (5).

Current data suggest that more than 20% of tumors contain mutations in GPCRs and G proteins (6). Activating mutations in GNAQ, GNA11 and GNAS have been shown to act as oncogenic drivers to promote tumorigenesis in several tumor types including uveal melanomas, intestinal cancers, and pancreatic adenocarcinomas (7–12). Previous work has also identified several activating mutations in the Gα subunits of the G_i/o_ subclass, specifically GNAI2 (R179C and R179H) and GNAO1 (R209C and R243H), in multiple tumor types including ovarian and adrenal tumors (13), as well as breast cancer and leukemia (14–16). A recent study by Song et al. has established the role of GNAO1 R209C in tumorigenesis of acute lymphoblastic leukemia (16). However, the functional significance of the other Gα_i/o_ mutants remains to be determined in clinically relevant tissues and models *in vivo.*

Our previous work found that many G_i/o_-coupled GPCRs are upregulated in breast cancer, and the upregulated G_i/o_-coupled GPCRs play a significant role in HER2-induced breast tumor initiation, growth, and metastasis (17, 18). G_i/o_-coupled GPCRs can transmit signals through both the dissociated Gα_i/o_ and Gβγ subunits. We and others previously showed that Gβγ serves as a point of convergence for many G_i/o_-coupled GPCR signals mediating breast cancer growth and metastasis (19, 20), but the role of Gα_i/o_ signaling in breast cancer progression remains to be characterized. In this study, we investigate the function of activating Gα_i/o_ mutations in breast cancer development, using cell lines and transgenic models. Our experiments revealed that the active Gα_i2_R179C and Gα_oA_R243H mutants alone were insufficient to induce breast cancer formation. In the transgenic mouse models of breast cancer induced by Neu expression or PTEN loss, the Gα_i2_R179C mutant enhanced spontaneous lung metastasis but did not affect mammary tumor formation and growth. Additional data showed that the active Gα_i2_R179C and Gα_oA_R243H mutants promote breast cancer metastasis through activation of the c-Src pathway. Together, our study not only elucidates the function of active GNAI2 and GNAO1 mutations in breast cancer metastasis but also provides compelling evidence to suggest that targeting the c-Src pathway may be an effective way to block tumor metastasis in breast cancer patients with active Gα_i/o_ mutants or elevated Gα_i/o_ signaling.

## Results

### The active Gα_i2_R179C and Gα_oA_R243H mutants alone are insufficient to induce mammary tumor formation

To determine the role of the Gα_i2_R179C and Gα_oA_R243H mutants in breast cancer formation, we expressed each of these mutants in a mammary epithelial cell line stably expressing firefly luminescence, MCF10A-FLU, using a doxycycline-inducible system (Figure 1A). As observed in Figure 1B, inducing expression of either Gα_i2_R179C or Gα_oA_R243H in MCF10A cells activated c-Src and STAT3 as previously reported for Gα_oA_R243H in NIH3T3 cells (15). Activation of both c-Src and STAT3 was successfully blocked using the specific inhibitors, saracatinib (Sara) and STAT3-IN-1 (STAT3i), respectively. Interestingly, inhibition of c-Src by saracatinib did not affect STAT3 activation by Gα_i2_R179C and Gα_oA_R243H. Similarly, inhibition of STAT3 by STAT3-IN-1 had no effect on c-Src activation (Figure 1B). These observations suggest that Gα_i2_R179C and Gα_oA_R243H activate c-Src and STAT3 through independent mechanisms.

**Figure 1.**
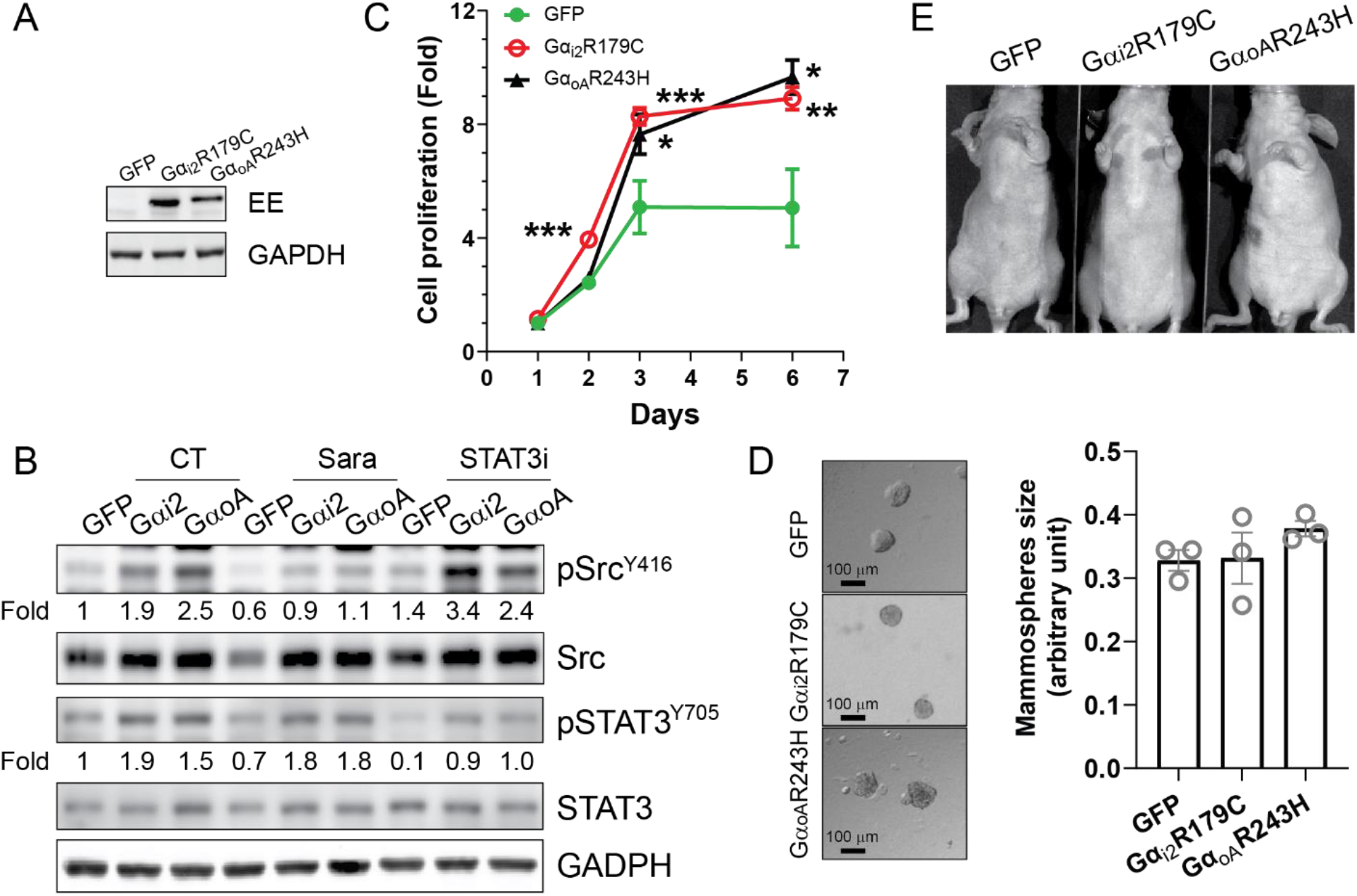
The active Gα_i/o_ mutants alone are insufficient to cause neoplastic transformation of mammary epithelial cells. **A-B**, representative immunoblots showing the expression of Gα_i2_R179C and Gα_oA_R243H in MCF10A cells (**A**) and activation of c-Src and STAT3 by phosphorylation of Src^Y416^ and STAT3^Y705^ in MCF10A cells treated with vehicle control (CT), 2 μM of saracatinib (sara) and STAT3-IN-1 (STAT3i) **(B)**. **CD**, the effect of Gα_i2_R179C and Gα_oA_R243H expression on the growth of MCF10A cells in 2D culture (**C**) and in Matrigel (**D**). **E**, representative images showing the absence of tumor formation in nude mice orthotopically implanted with MCF10A cells expressing GFP, Gα_i2_R179C or Gα_oA_R243H. *, **,*** p<0.05, 0.01 and 0.001 *vs.* GFP, n=4-6. One-way ANOVA was used for statistical analysis in this Figure.

In 2D cell culture, MCF10A cells expressing Gα_i2_R179C or Gα_oA_R243H grew faster than control cells expressing GFP (Figure 1C). However, when grown in Matrigel, these cells did not show a significant difference in growth (Figure 1D). We then implanted MCF10A cells expressing either GFP, Gα_i2_R179C, or Gα_oA_R243H into the mammary gland of nude mice. After monitoring the mice for tumor growth over a period of four months, using mammary palpation and highly sensitive bioluminescence imaging, we found that none of the cells formed detectable tumors in any of the mice (Figure 1E). These results suggest that activation of Gα_i/o_ signaling alone is insufficient to drive the formation of breast tumors.

To further confirm these findings, we generated mice to selectively express Gα_i2_R179C in the mammary gland using the tetO promoter system. We crossed transgenic mice carrying Gα_i2_R179C under the tetO promoter with transgenic mice expressing the transactivator, tTA, from the mammary gland-specific MMTV promoter, MMTV-tTA. Western blot comparison of mammary glands from 5-month-old mice expressing tTA alone and tTA/ Gα_i2_R179C (Gα_i2_R179C) confirmed Gα_i2_R179C expression in the tTA/Gα_i2_R179C mice in the absence of doxycycline treatment (Figure 2A). Neither the littermate control (tTA) mice, nor Gα_i2_R179C mice formed palpable tumors for up to 20 months. Additionally, whole-mount in situ staining of mammary glands extracted from 3- and 13-month-old tTA and Gα_i2_R179C 2 mice did not reveal any differences in morphology (Figure 2B). This indicates that Gα_i2_R179C does not affect normal mammary development and does not induce mammary gland hyperplasia. Together, these results indicate that activating mutations in GNAI2 or GNAO1 alone are insufficient to induce breast cancer formation.

**Figure 2.**
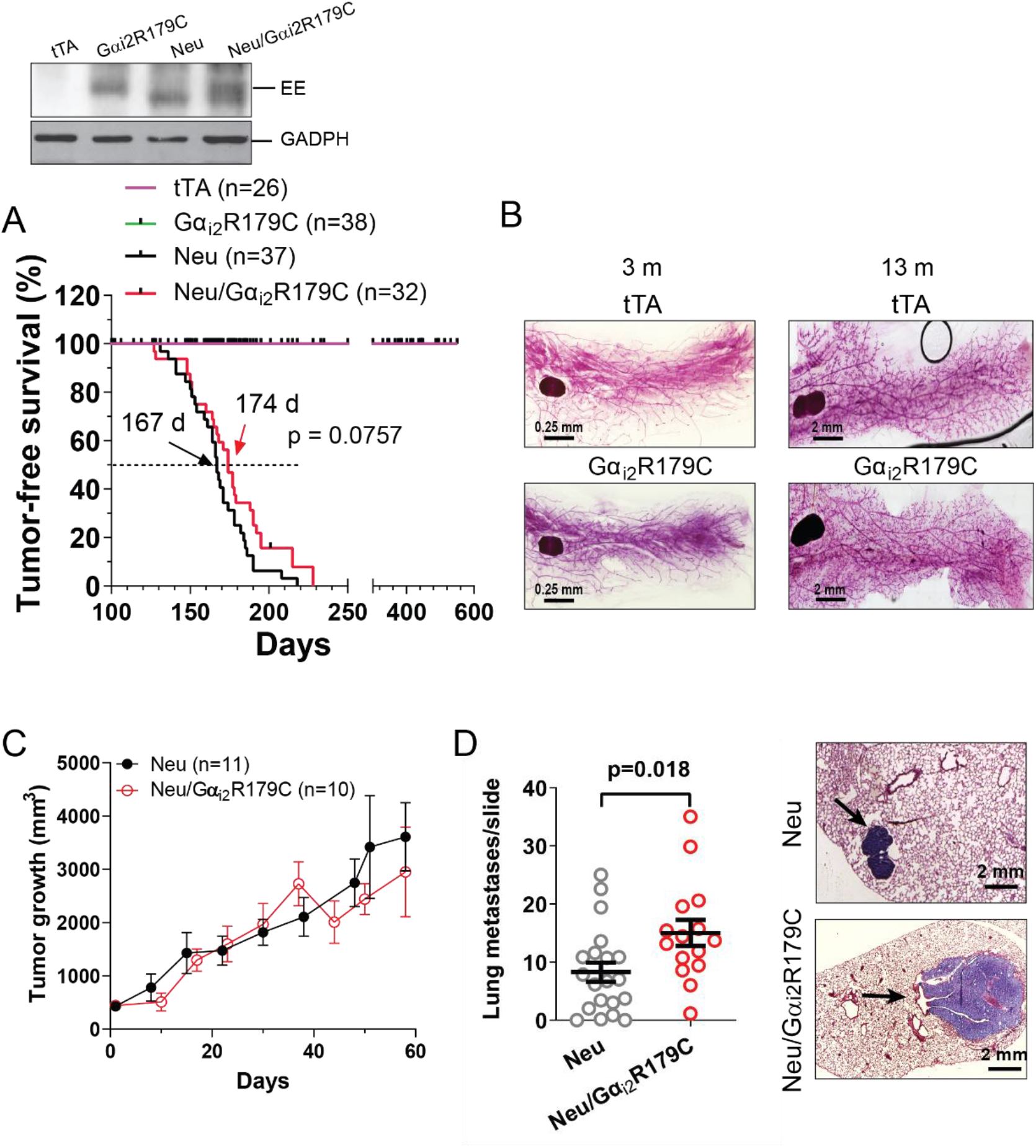
Expression of G_αi_2R179C neither induces mammary tumor formation nor affects HER2-driven primary tumor development, but accelerates lung metastasis. **A**, tumor-free survival curves of mice expressing tTA, tTA/Gα_i2_R179C (Gα_i2_R179C), tTA/Neu (Neu), and tTA/Neu/Gα_i2_R179C (Neu/ Gα_i2_R179C), and representative immunblots showing ee-tagged Gα_i2_R179C expression in the mammary gland and Neu tumors. **B**, whole-mount *in situ* staining showing the morphology of the mammary glands from 3 and 13-month-old tTA and Gα_i2_R179C mice. **C**, Neu and Neu/Gα_i2_R179C tumor growth curves. **D**, a graph to show the number of lung metastases and images representative of HE-stained lung metastasis (indicated by arrows) from the transgenic Neu and Neu/Gα_i2_R179C mice. Two tail unpaired Student’s *t* test was used for statistical analysis in this Figure.

### Gα_i/o_ signaling promotes breast cancer metastasis

Since G_i/o_-GPCR signaling is upregulated in HER2^+^ breast cancer, we further explored the role of elevated Gα_i/o_ signaling in Neu tumor formation. We generated trigenic mice, tTA/Gα_i2_R179C/Neu (Neu/ Gα_i2_R179C), by crossing tTA/Gα_i2_R179C mice with MMTV-Neu (Neu) transgenic mice, which selectively express an activated rat ErbB2/HER2 homologue in their mammary glands. The Neu mice spontaneously formed mammary tumors, which then metastasized to the lungs. The median time for tumor formation in the Neu mice (167 days) was not significant different from the Neu/ Gα_i2_R179C mice (174 days) (Figure 2A). There was also no significantly difference between tumor growth in Neu mice compared to Neu/ Gα_i2_R179C mice (Figure 2C). However, Gα_i2_R179C significantly increased the number of Neu-induced lung metastases (Figure 2D).

To test whether the Gα_i2_R179C mutant might regulate the progression of breast cancer that has already been induced by other existing genetic abnormalities, such as deletion of the tumor suppressor gene, PTEN (Phosphatase And Tensin Homolog), we crossed tTA/Gα_i2_R179C mice with transgenic mice exhibiting mammary gland-specific deletion of one PTEN allele (MMTV-Cre/PTEN^fl/+^; PTEN^+/-^). As seen in Figure 3A, PTEN^+/-^ mice generated mammary tumors after an extended latency (407 days) compared to Neu mice (167 days). Following a similar trend as we saw previously with Neu/ Gα_i2_R179C mice, Gα_i2_R179C expression in PTEN^+/-^ mice (PTEN^+/-^ /Gα_i2_R179C) did not affect PTEN loss-induced tumor onset and growth (Figure 3A and 3B), but it did significantly increase the number of lung metastases (Figures 3C and 3D). These results suggest that although Gα_i_ signaling is not required for the primary growth of mammary tumors, it is critical in the metastasis of tumors to distant organs.

**Figure 3.**
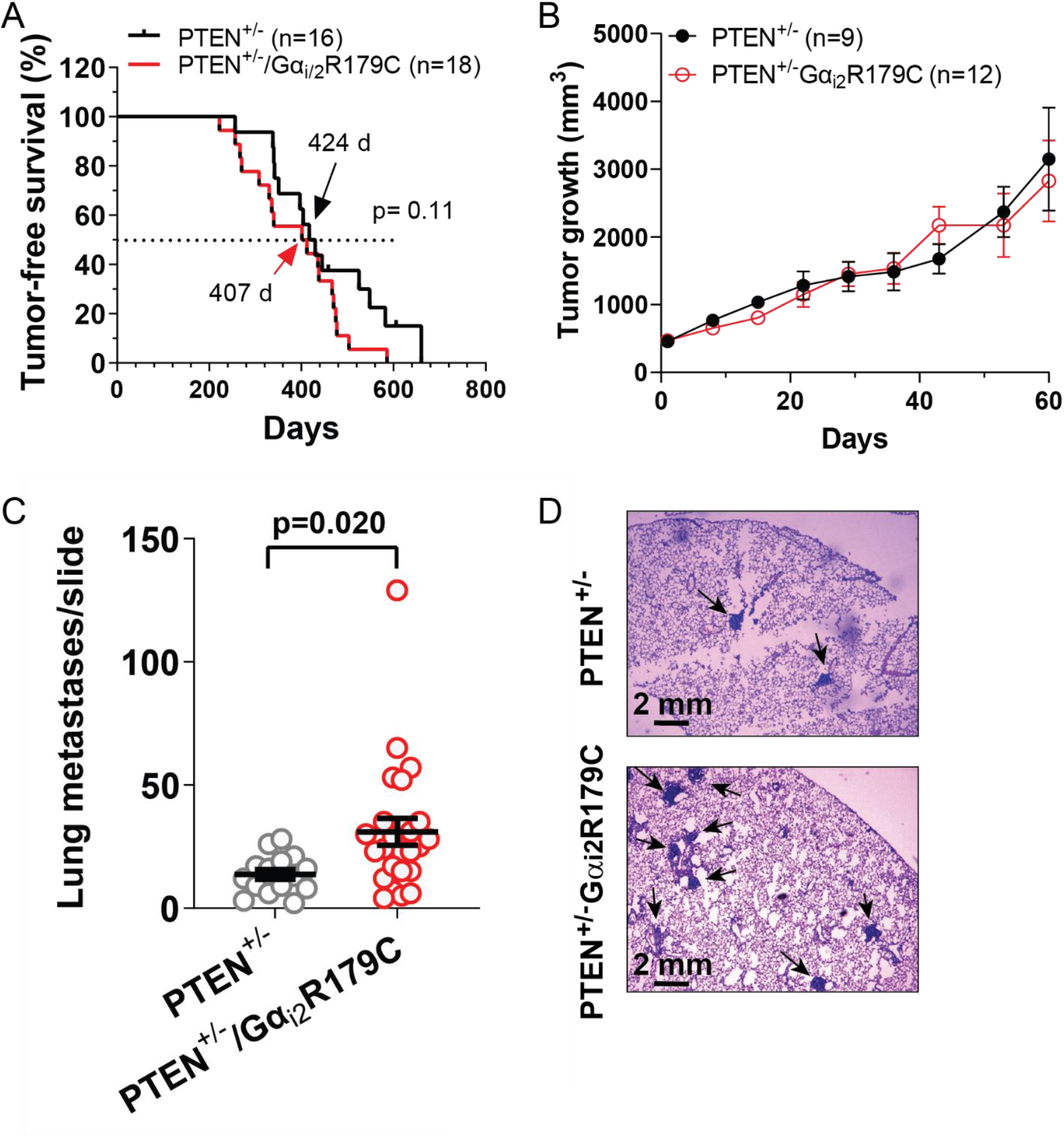
Expression of Gα_i2_R179C enhanced PTEN deficiency-driven lung metastasis. **A**, tumor-free survival curves of MMTV-Cre/PTEN^fl/+^ (PTEN^+/-^) and MMTV-Cre/PTEN^fl/+^/MMTV-tTA/Gα_i2_R179C (PTEN^+/-^/Gα_i2_R179C) mice. **B**, PTEN^+/-^ and PTEN^+/-^/Gα_i2_R179C tumor growth curves. **C-D**, a graph to show the number of lung metastases **(C)** and images representative of HE-stained lung metastases (indicated by arrows) **(D)** from PTEN^+/-^ and PTEN^+/-^/Gα_i2_R179C mice. Two tail unpaired Student’s *t* test was used for statistical analysis in this Figure.

We then question whether Gα_oA_ signaling may play a similar role as Gαi signaling in promoting breast tumor metastasis. To this end, we injected Neu cells expressing inducible GFP or Gα_oA_R243H into the left ventricle of nude mice to mimic the metastatic spread of tumor cells into multiple organs. Beginning immediately after injection, the mice were continuously fed with doxycycline-containing chow to induce GFP and Gα_oA_R243H expression. Bioluminescence imaging showed that the Gα_oA_R243H-injected mice exhibited enhanced tumor growth compared to the GFP-injected mice (Figure 4A). While all the injected mice formed tumors in multiple organs, in particular the lung, there were significantly more metastatic tumors found in the mice injected with Gα_oA_R243H-expressing Neu cells (Figure 4B). These results therefore confirm that Gα_oA_ signaling is also important in tumor metastasis.

**Figure 4.**
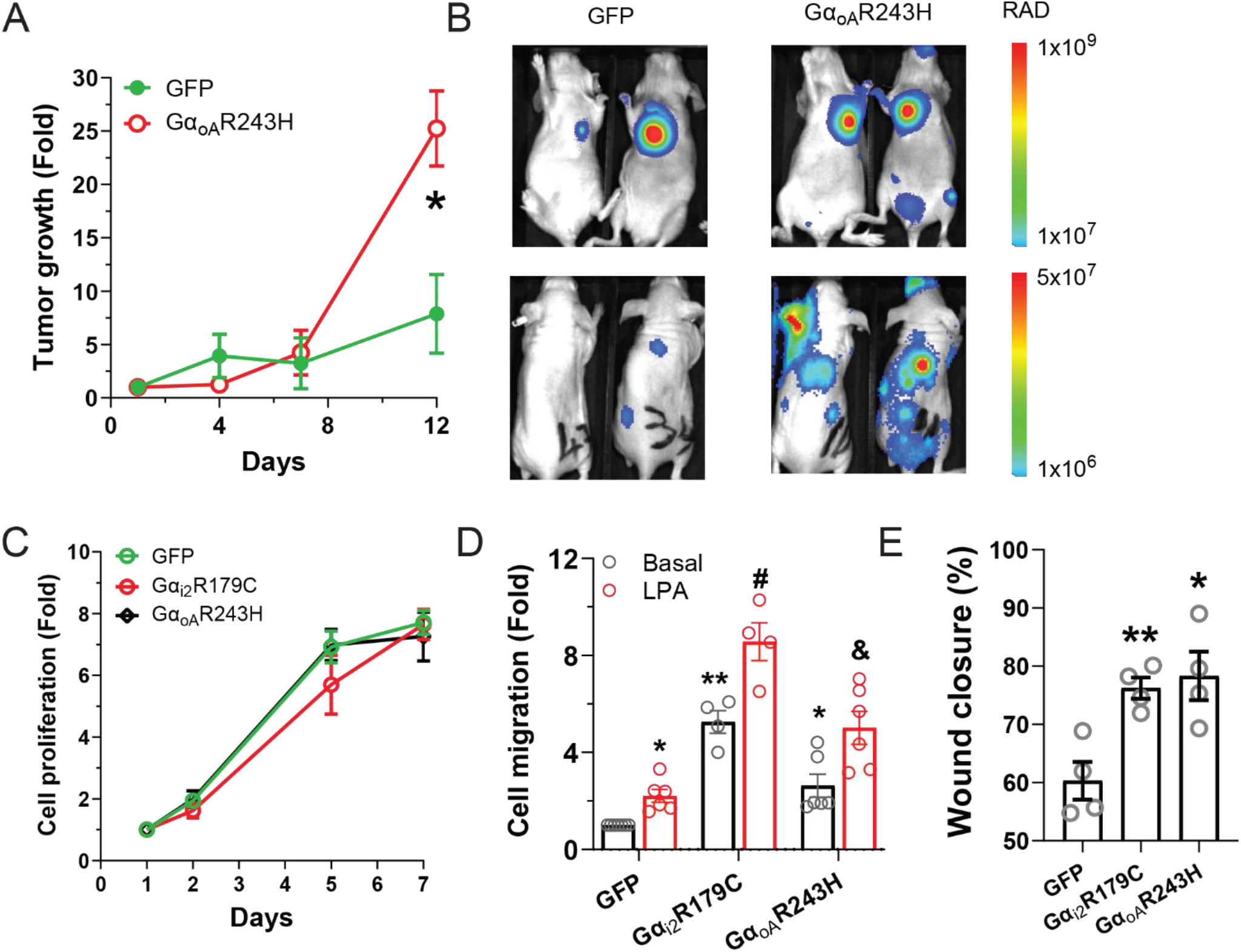
Effects of Gα_i2_R179C and Gα_oA_R243H on Neu cell growth and migration and tumor metastasis. **A-B**, the tumor growth curves of Neu cells expressing GFP or Gα_oA_R243H injected into nude mice via left ventricle (**A**) and representative bioluminescence images of mice at the dorsal (top) and ventral (bottom) positions (**B**). **C**, the *in vitro* growth curves of Neu cells expressing GFP, Gα_i2_R179C, or Gα_oA_R243H. **D,** basal and LPA (10 μM)-stimulated transwell migration of Neu cells expressing GFP, Gα_i2_R179C, or Gα_oA_R243H. **E,** wound healing assay data of Neu cells expressing GFP, Gα_i2_R179C, or Gα_oA_R243H. *, **p<0.05 and 0.01 *vs* GFP basal; ^#, &^ p<0.05 and 0.01 *vs* GFP LPA. Two-way and one-way ANOVA were used for statistical analysis in Figure D and E, respectively.

To understand the mechanisms by which Gα_i_ and Gα_oA_ signaling promote tumor metastasis, we determined their impact on cell proliferation and migration. Neu cells expressing GFP, Gα_i2_R179C, and Gα_oA_R243H grew at a similar rate, showing no significant difference in proliferation capacity (Figure 4C). Notably, however, the Neu cells expressing Gα_i2_R179C or Gα_oA_R243H exhibited enhanced trans-well migration ability compared to cells expressing GFP (Figure 4D). This difference is evident in the basal state but is amplified upon treatment with LPA. Data from wound-healing assays also demonstrated increased migration capacity of Gα_i2_R179C and Gα_oA_R243H expressing cells compared to cells expressing GFP (Figure 4E). These results suggest that Gα_i/o_ signaling may promote tumor metastasis by enhancing tumor cell motility.

### Gα_i/o_ signaling promotes tumor metastasis via the c-Src pathway

We showed previously that G_i/o_-coupled GPCRs transactivate EGFR and HER2, and that their signaling converges at the AKT and c-Src pathways to promote Neu-induced breast cancer growth and metastasis (17). Western blotting analysis shows that, compared with Neu tumors, Gα_i2_R179C tumors showed significantly increased phosphorylation of EGFR^Y1068^ and c-Src^Y416^ but no significant changes in phosphorylation of HER2^Y1221/1222^, STAT3^Y705^ and AKT^S473^ (Figure 5A and 5B). Similarly, phosphorylation of c-Src^Y416^, but not AKT^S472^, STAT3^Y705^ and EGFR^Y1068^, was significantly increased in PTEN^+/-^ /Gα_i2_R179C tumors as compared to PTEN^+/-^ tumors (supplemental Figure 1A and 1B). These findings suggest that Gα_i/o_ signaling might be involved in G_i/o_-GPCR-stimulated c-Src activation, and in a context-dependent manner, EGFR transactivation.

**Figure 5.**
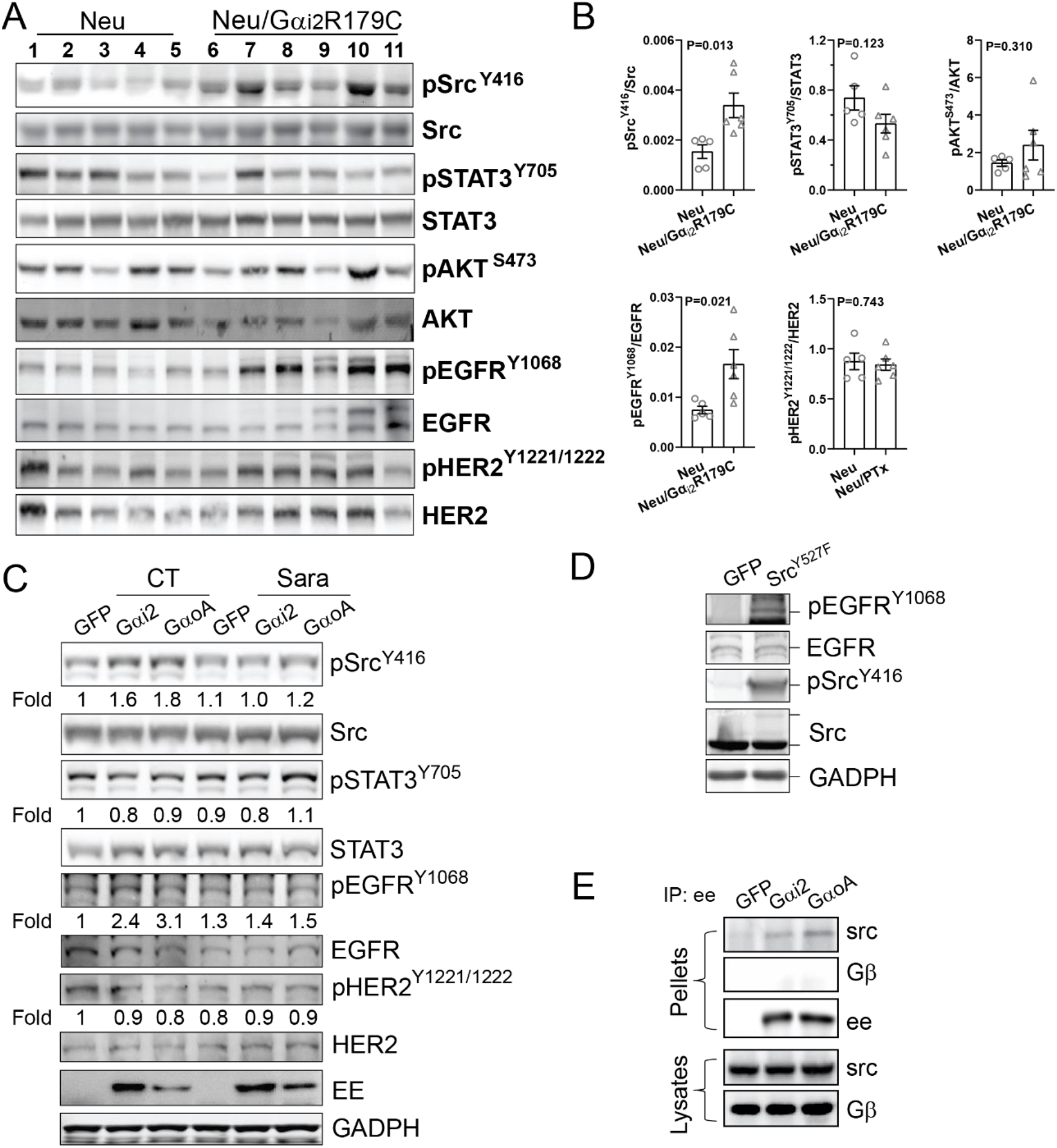
Gα_i2_R179C and Gα_oA_R243H activate c-Src and induce EGFR transactivation in tumor cells. **A-B**, Western blotting (**A**) showing increased phosphorylation of c-Src^Y416^ and EGFR^Y1068^ and no change in phosphorylation of AKT^s473^, STAT3^Y705^ and HER2^Y1221/1222^ in Neu/Gα_i2_R179C tumors as compared to Neu tumors. Each lane represents a sample from an individual tumor. The Western blotting data from **A** were quantified and expressed as the ratio of the phosphorylated to total proteins (**B**). Two tail unpaired Student’s *t*test was used for statistical analysis of the data in **B**, and p values are shown. **C-D**, Western blotting showing phosphorylation of the indicated proteins in Neu cells expressing GFP, Gα_i2_R179C (Gα_i2_) or Gα_oA_R243H (Gα_oA_) and treated with vehicle control (CT) or saracatinib (sara; 2 μM) (**C**) and Neu cells overexpressing GFP or the constitutively active c-Src^Y527F^ mutant (**D)**. **E**, co-immunoprecipitation of ee-tagged Gα_i2_R179C and Gα_oA_R243H with c-Src in Neu cells. Neu cells expressing GFP, Gα_i2_R179C (Gα_i2_) or Gα_oA_R243H (Gα_oA_) were immunoprecipitated with the anti-ee antibody and probed with the indicated antibodies.

Additional immunoblotting data showed that phosphorylation of EGFR^Y1068^ and c-Src^Y416^ was increased in Neu cells expressing Gα_i2_R179C or Gα_oA_R243H, as compared to Neu cells expressing GFP (Figure 5C). In contrast, there appears to be no significant difference in AKT^S473^, STAT3^Y705^ and HER2^Y1221/1222^ phosphorylation (Figure 5C). Furthermore, treatment of Neu cells with the Src-specific inhibitor, saracatinib, abolished the observed increase in both c-Src^Y416^ and EGFR^Y1068^ phosphorylation in Neu cells expressing Gα_i2_R179C or Gα_oA_R243H. This finding suggests that EGFR is likely a downstream target for transactivation by c-Src. This proposition was corroborated by the finding that EGFR^Y1068^ phosphorylation was increased after inducing expression of a constitutively active c-Src mutant, c-Src^Y527F^ in Neu cells (Figure 5D).

We then sought to identify the mechanisms by which Gα_i2_R179C and Gα_oA_R243H may activate c-Src. We first considered the possibility of the Gβγsubunit being involved. Theoretically, the active Gα_i/o_ mutants could release free Gβγ subunits, which could activate c-Src independent of the Gα_i/o_ subunits (21). To test this, we inhibited Gβγ signaling by overexpressing Gα_t_, a scavenger of Gβγ, in Neu cells as we previously reported (20). As expected, overexpression of Gα_t_ suppressed LPA-stimulated ERK phosphorylation, which is known to be activated by Gβγ (20), in Neu cells expressing GFP or Gα_i2_R179C (supplemental Figure 2A). However, the enhanced c-Src activation in Neu cells expressing Gα_i2_R179C or Gα_oA_R243H was unaffected by Gα_t_ overexpression (supplemental Figure 2B). These findings indicate that the active Gα_i2_R179C and Gα_oA_R243H mutants activate c-Src independent of the dissociated Gβγ subunits.

We then pursued the idea that Gα_i2_R179C and Gα_oA_R243H may activate c-Src by direct interaction as previously reported for Gα_i_ (22). We performed co-immunoprecipitation assays with Neu cell lysates and found that Gα_i2_R179C and Gα_oA_R243H interacted with c-Src, but not Gβγ (Figure 5E). Transwell migration assays also showed that overexpression of Gα_t_ or G protein coupled receptor kinase 2 (GRK2-ct), another Gβγ scavenger, did not affect the enhanced Neu cell migration induced by Gα_oA_R243H (supplemental Figure 2C). Suppression of c-Src by saracatinib, however, abolished the enhanced cell migration by Gα_i2_R179C and Gα_oA_R243H while not significantly altering the migration of GFP-expressing cells (Figure 6A). These results indicate that Gα_i2_R179C and Gα_oA_R243H promote tumor cell migration via the c-Src pathway.

**Figure 6.**
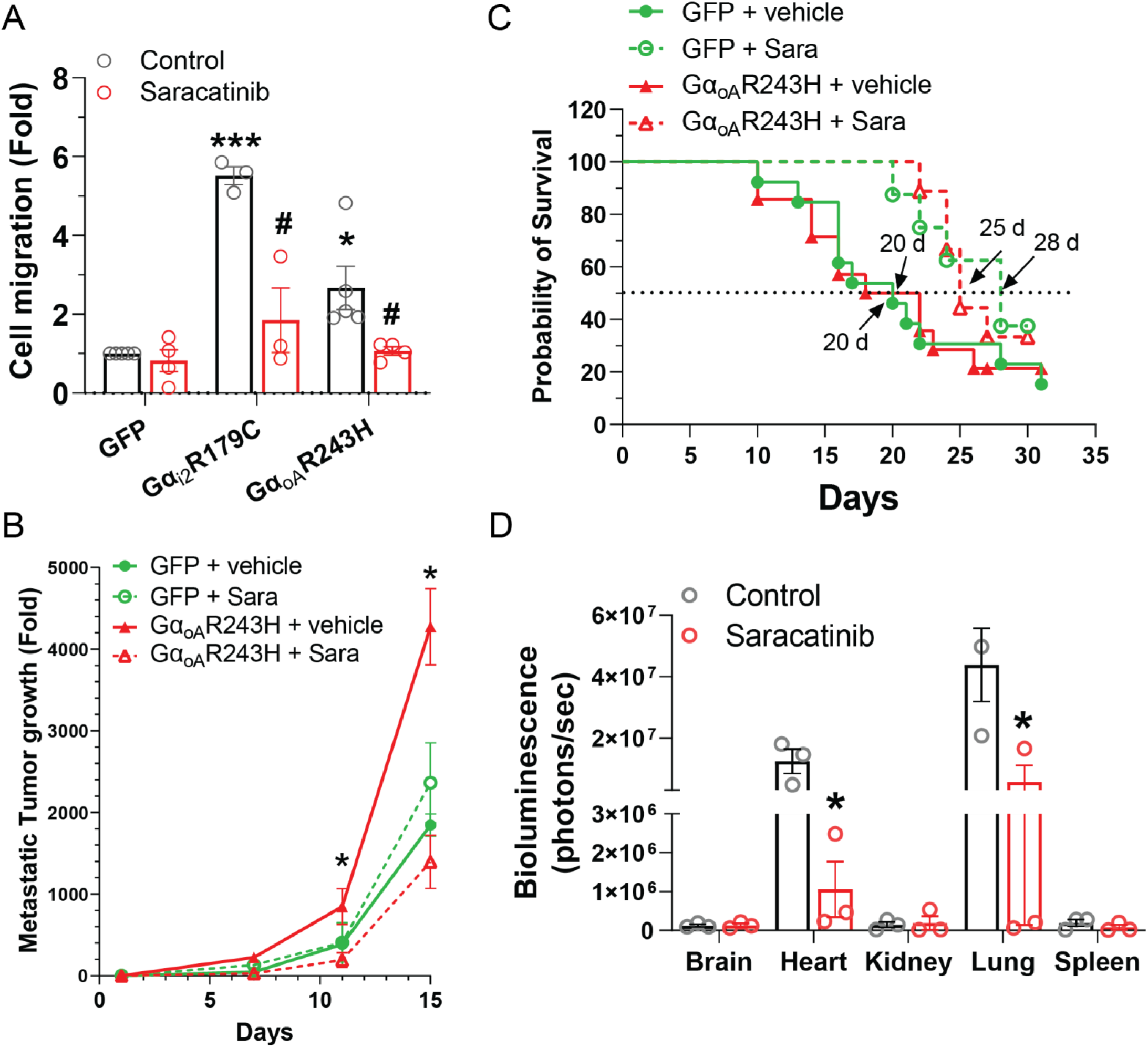
The effects of saracatinib on Neu cell migration and tumor metastasis. **A**, transwell migration of Neu cells expressing GFP, Gα_i2_R179C or Gα_oA_R243H and treated with vehicle (control) or saracatinib (2 μM). *, ***p<0.05 and 0.001 *vs* GFP control; #p<0.05 *vs* Gα_i2_R179C or Gα_oA_R243H control. **B-C**, the growth curves (**B**) and survival curves (**C)** of mice harvesting GFP and Gα_oA_R243H metastatic tumors and treated with vehicle or saracatinib (25mg/kg initially for 7 days and then 50mg/kg for the rest of treatment, gavage daily; sara). Two-way ANOVA was used for statistical analysis in Figure B and C (n=9-13). Log-rank test was used for statistical analysis in Figure D (n=9-13). **D**, a graph showing the bioluminescence intensity of the indicated tissues measured via *ex vivo* bioluminescence imaging from the remaining mice harvesting GFP metastatic tumors and treated with vehicle (control) or saracatinib for two weeks. *p<0.05 vs vehicle.

To validate our suppositions *in vivo*, we established tumor metastases in multiple organs by injecting Neu cells expressing GFP or Gα_oA_R243H into the left ventricle of nude mice and feeding them doxycycline-containing chow to induce GFP and Gα_oA_R243H expression. Eight days post-injection, the mice were treated with a vehicle control or saracatinib and tumor growth was monitored by bioluminescence imaging. As measured by whole-body bioluminescence intensity, saracatinib treatment did not affect the growth of GFP-expressing Neu tumors but diminished the enhanced growth of Gα_oA_R243H-expressing Neu tumors (Figure 6B. Moreover, the overall survival of mice injected with either GFP- or Gα_oA_R243H-expressing Neu cells was significantly increased by saracatinib (Figure 6C; 20 and 20 days for vehicle control *vs* 28 and 25 days for saracatinib, respectively). At the end of the treatment for two weeks, *ex vivo* bioluminescence imaging of key tissues from the mice harvesting GFP tumors showed that GFP tumors primarily grew in the lung and heart and saracatinib treatment significantly reduced tumor growth in these tissues (Figure 6D). Together, these results indicate that Gα_i/o_ signaling can accelerate Neu tumor metastasis via the c-Src pathway.

## Discussion

In this study, we demonstrated that overexpression of the constitutively active Gα_i2_R179C or Gα_oA_R243H mutant alone in mammary epithelial cells is insufficient to induce mammary tumor formation but can cooperate with Neu/HER2 hyperactivation or PTEN deficiency to drive tumor metastasis. Somatic GNAI2 R179C and GNAO1 R243H mutations have been found in human cancers, including ovarian and adrenal tumors as well as breast cancer (13–15). Although these mutants were shown to promote oncogenic transformation and anchorage-independent growth of fibroblasts or mammary epithelia cells *in vitro,* their function as oncogenes has never been characterized in relevant tissues *in vivo*. Thus, our study provides the first evidence for the significant role of these mutants in breast cancer progression.

Notably, the active Gα_i2_R179C and Gα_oA_R243H mutants selectively affect breast tumor cell migration and tumor metastasis but have no effect on tumor cell proliferation *in vitro*. The Gα_i2_R179C mutant also shows no significant effect on tumor initiation and growth in the two different genetic mouse models of breast cancer, Neu overexpression and PTEN deficiency. The mechanisms by which Gα_i2_R179C and Gα_oA_R243H selectively enhance tumor cell migration and tumor metastasis are unclear but may be related to their ability to activate only a limited number of pathways. Indeed, in Neu cells and tumor samples, Gα_i2_R179C and Gα_oA_R243H primarily enhance c-Src and EGFR transactivation. This is in stark contrast to our previous data showing that G_i/o_-coupled receptor signaling contributes to the activation of multiple pathways, including c-Src and AKT, and transactivation of both EGFR and HER2 (17). The activation of additional pathways by G_i/o_-GPCRs likely involve Gβγ subunits liberated from heterotrimeric G_i/o_ proteins since activation of heterotrimeric G_i/o_ proteins by GPCRs generates both functional Gα_i/o_ and Gβγ subunits and Gβγ is known to be involved in PI3K/AKT activation (3, 4, 23–25). Moreover, we and others have established that Gβγ signaling is critical for breast cancer cell growth and migration induced by multiple G_i/o_-GPCRs (20, 24). Given that blocking G_i/o_-coupled receptor signaling not only delays Neu-induced tumor initiation, but also suppresses tumor growth and metastasis (17), G_i/o_-coupled GPCRs likely signal through both Gα_i/o_ and Gβγ to accelerate breast tumor progression. While Gα_i/o_ signaling primarily drives tumor metastasis, Gβγ signaling may promote both primary tumor growth and metastasis.

Our data indicate that Gα_i2_R179C and Gα_oA_R243H primarily activate the c-Src pathway to promote tumor metastasis. In breast tumor cells and tissues, overexpression of Gα_i2_R179C and Gα_oA_R243H enhanced c-Src activation. These findings are consistent with several reports that active Gα_oA_ mutants can activate c-Src in fibroblasts (15, 26). The activation of c-Src by Gα_i2_R179C and Gα_oA_R243H does not seem to involve Gβγ since co-expression with a Gβγ scavenger has no effect on c-Src activation. Rather, Gα_i2_R179C and Gα_oA_R243H may activate c-Src by direct interaction as previously reported for active Gα_i_ since they are co-immunoprecipitated with c-Src in breast tumor cells (22). We cannot, however, exclude the possibility that there are other proteins mediating or regulating this interaction.

The activation of c-Src by Gα_i2_R179C and Gα_oA_R243H is functionally important since blocking c-Src activation with a specific inhibitor, saracatinib, suppresses the mutant-promoted tumor cell migration *in vitro* and metastatic spread of tumor cells to multiple organs *in vivo.* Saracatinib, however, has no significant effect on Neu-driven tumor cell migration and tumor metastasis, although it significantly improves the survival of Neu mice. The reason for the differential effect of saracatinib on GFP tumor growth and overall survival is unclear but may be related to the inhibition of metastatic tumor growth in critical organs such as the lung and heart, as indicated by *ex vivo* bioluminescence imaging of the tissues from the surviving mice at the end of the experiments. These results suggest that Neu-driven tumor metastasis may involve multiple redundant signaling pathways and combining Src inhibition with other therapeutic reagents may be required for effectively blocking Neu tumor metastasis.

We also found that activation of c-Src by Gα_i2_R179C and Gα_oA_R243H causes EGFR transactivation in Neu cells and Neu tumors, supporting a functional interaction between c-Src and EGFR as previously reported (27). However, although Gα_i2_R179C also enhances c-Src activation in PTEN tumors, it does not cause EGFR transactivation, suggesting the role of c-Src in EGFR transactivation is context dependent. Since Gα_i2_R179C promotes tumor metastasis in both Neu and PTEN tumor models, these results also suggest that the transactivation of EGFR by c-Src is not essential for Gα_i2_R179C- and Gα_oA_R243H-promoted tumor metastasis.

Once activated, c-Src can interact with various substrates and effectors, including STAT3, to promote tumor cell proliferation, migration, and invasion (28, 29). Previous studies showed that the active Gα_i2_ and Gα_oA_ mutants can induce oncogenic transformaton of NIH3T3 fibroblasts via the c-Src-stimulated STAT3 pathway (26, 30). We found that Gα_i2_R179C and Gα_oA_R243H can activate STAT3 in MCF10A cells but have no effects in Neu cells. Moreover, the expression of Gα_i2_R179C enhances c-Src but not STAT3 activation in Neu and PTEN^+/-^ tumors. The activation of STAT3 by Gα_i2_R179C and Gα_oA_R243H in MCF10A cells also does not seem to involve c-Src. These findings suggest that the activation of STAT3 by the Gα_i2_ and Gα_oA_ mutants is likely to be cell type dependent. Further studies are required to identify the downstream effectors of c-Src that mediate Gα_i2_R179C- and Gα_oA_R243H-promoted tumor cell migration and metastasis.

In conclusion, we demonstrate through both *in vitro* and *in vivo* studies that active Gα_i/o_ mutants accelerate breast cancer metastasis via the c-Src pathway. In addition to mutations in Gα_i/o_ genes, multiple G_i/o_-coupled receptors and Gα_i/o_ proteins are frequently upregulated in many cancers including breast cancer (6, 17, 31). Moreover, the activity of c-Src is increased in a variety of neoplastic tissues and targeting c-Src has been proposed as a promising strategy for blocking breast cancer metastasis (29, 32–34). Given that tumor metastasis is a major cause of cancer mortality, our findings that c-Src is a key mediator of Gα_i/o_ signaling in promoting tumor metastasis have important ramifications in the development of personalized therapies to improve the survival of breast cancer patients with active Gα_i/o_ mutants or enhanced Gα_i/o_ signaling.

## Methods

### Reagents

Saracatinib was obtained from LC Laboratories. Antibodies for EGFR (no. 2232), phospho-EGFR^Y1068^(no. 3777), EGFR (no. 4267), HER2 (no. 2165), phospho-HER2^Y1221/1222^ (no. 2243), AKT (no. 4685), phospho-AKT^S473^ (no. 4060), Src (no. 2109), phospho-Src^Y416^ (no. 6943), ERK1/2 (no. 4696) and phospho-ERK1/2^T202/Y204^ (no. 4370) were obtained from Cell Signaling Technology; GADPH (sc-47724) from Santa Cruz Biotechnology; mouse anti-ee antibody (no. 901801) from Biolegend.

### Cell lines

Neu cells were generated from tumors formed by the transgenic mice, MMTV-c-Neu, and cultured in DMEM media containing 10% FCS. Cell lines were tested for *Mycoplasma* using a Mycoplasma Detection kit (ATCC). MCF10A cells were purchased from the ATCC, and cultured in DMEM/F12 media containing 5% horse serum supplemented with EGF at 20ng/ml, hydrocortisone at 500ng/ml, cholera toxin at 100ng/ml, and insulin at 10μg/ml. Each cell line was cryopreserved at low passage numbers (less than six passages after receipt) and used in experiments for a maximum of 18 passages.

### Plasmid

The lentiviral vector (Tet-CA-Src-GFP) for tetracycline-inducible expression of the GFP-tagged, constitutively active Src mutant, Src/Y527F, was obtained from Addgene (item #83469). Gαi2R179C and Gα_oA_R243H mutants were constructed using pcDNA3.1 wild-type human ee-tagged-Gα_i2_- and ee-tagged-Gα_oA_ plasmids as template (cDNA Resource Center) and Q5 Site-directed mutagenesis kit (New England BioLabs). The lentiviral vector for tetracycline-inducible expression of the Gα_i2_R179C and Gα_oA_R243H mutants was constructed by first cloning Gα_i2_R179C and Gα_oA_R243H from pcDNA3.1 to pENTR vector (Thermofisher Scientific) and then into the destination vector pLIX_402 (Addgene, item #41394) using the Gateway cloning system.

The adenovirus vectors for Gα_t_ and GRK2ct were constructed by first cloning Gα_t_ and GRK2ct into the pENTR vector and then into the destination vector, pAd/CMV/DEST (Thermofisher Scientific), using the Gateway cloning system.

### Lentiviral production

Lentiviruses were generated in HEK293FT cells as described previously (23). Lentiviruses collected from the cell culture supernatants were concentrated using the Lenti-X-concentrator (Takara Bio).

### Adenoviral production and transduction

Adenoviruses were generated in HEK293A cells using the ViraPower Adenoviral Expression System (Thermofisher Scientific) and purified with a CsCl_2_ density gradient combined with ultracentrifugation. Transduction of Neu cells was conducted by incubation of cells with purified adenoviruses overnight.

### Establishment of stable cell lines

The MCF10A and Neu cells were transduced with lentiviruses encoding GFP-tagged Src^Y527F^, GFP, Gα_i2_R179C, or Gα_oA_R243H, and selected with puromycin (2 μg/ml) for at least 1 week to establish stable lines.

### Cell proliferation and viability assays

Cell proliferation in two-dimensional culture or in Matrigel was analyzed as we described previously (17, 23). Cell viability was quantified using AlamarBlue (Thermo Fisher Scientific) assays as per the manufacturer’s instructions.

### Cell migration and wound healing assays

Transwell migration and wound healing assays were performed as we described previously (17, 23). To exclude the influence of cell proliferation, cells were treated with 5 μg/ml mitomycin.

### Western blotting analysis

Protein lysates were prepared from cells and tumor tissues and analyzed by Western blotting as we described. Blots were imaged using the iBright 1500 (Thermo Fisher Scientific) or Odyssey (LI-COR Biotechnology) imaging system (17, 23).

### Co-immunoprecipitation assay

Neu cells were lysed in modified RIPA buffer containing 50mM Tris-HCl, 150mM NaCl, 1mM EDTA and 1% Triton, pH 7.4 as we reported previously. Cell lysates were incubated overnight at 4 °C with mouse anti-ee antibodies pre-bound to Dynabeads protein G (Thermo Fisher Scientific(. The immunoprecipitated protein complex was analyzed by Western blotting.

### Mouse studies

MMTV-c-Neu (no. 005038), TetO-Gα_i2_R179C (no. 017333), MMTV-Cre (n0. 003553) and PTEN^fl/fl^ (no. 034621) mice were purchased from the Jackson Laboratory and MMTV-tTA mice were obtained from Dr. Wagner’s laboratory (50). All mice were in the FVB/N genetic background. Mice were genotyped by PCR as reported previously (24, 50, 51). Female mice were kept as virgins throughout the experiments. To determine tumor onset, mice were checked twice per week by palpation beginning four months after birth. To assess tumor progression, the largest tumor was measured weekly by caliper. To evaluate lung metastasis, the lung was harvested once the largest tumor reached a size of 2 cm in diameter and was perfused and fixed with 4% paraformaldehyde before paraffin embedding. The number of metastases in the lung was analyzed by serial sectioning followed by HE staining.

To generate metastatic mouse models, Neu cells (~5x 10^5^ in 100 μl of phosphate-buffered saline) expressing luciferase and doxycycline-inducible GFP or Gα_oA_R243H were injected into 6–8-week-old female nude mice (Charles River) via left ventricle. Immediately post-injection, bioluminescence imaging was performed to confirm successful injection. Mice were then continuously fed with a doxycycline-containing diet (TD.01306; ENVIGO) to induce target gene expression. Tumor growth was monitored by bioluminescence imaging of the mice at both dorsal and ventral positions as we reported previously. To determine the effect of c-Src inhibition, mice were administered with saracatinib (25mg/kg, daily gavage) dissolved in 2% DMSO, 30% PEG300 and H_2_O, eight days post-injection.

### Statistics

Data were expressed as mean ± SEM. Statistical comparisons between groups were analyzed by two tail Student’s *t* test or ANOVA (*P*<0.05 was considered significant). The survival curves were analyzed according to the Kaplan–Meier method.

## Study approval

All animal studies were conducted in accordance with an IACUC-approved protocol at the University of Iowa.

## Authors’ Contributions

Data curation, C. Lyu, A. Bhimani and W. Draus. Formal analysis, C. Lyu and S. Chen; Methodology, C. Lyu; Project administration, R. Weigel and S. Chen; Writing-original draft, S. Chen.; Writing-review & editing, A. Bhimani, R. Weigel and S. Chen.

## Acknowledgements

We would like to thank Dr. Yuanchao Ye and Maddison Lensing for their assistance in some of animal breeding and studies. This work was supported in part by NIH grant R01CA207889 and DOD BCRP breakthrough award level 2 (BC151478), and an NIH shared instrumentation grant (1S10OD026835-01).

## References

1. Bhushan A, Gonsalves A, and Menon JU. Current State of Breast Cancer Diagnosis, Treatment, and Theranostics. Pharmaceutics. 2021;13(5).

2. Campbell AP, and Smrcka AV. Targeting G protein-coupled receptor signalling by blocking G proteins. Nat Rev Drug Discov. 2018;17(11):789–803.

3. Oldham WM, and Hamm HE. Heterotrimeric G protein activation by G-protein-coupled receptors. Nat Rev Mol Cell Biol. 2008;9(1):60–71.

4. Smrcka AV. Molecular targeting of Galpha and Gbetagamma subunits: a potential approach for cancer therapeutics. Trends in pharmacological sciences. 2013;34(5):290–8.

5. Arang N, and Gutkind JS. G Protein-Coupled receptors and heterotrimeric G proteins as cancer drivers. FEBS Lett. 2020;594(24):4201–32.

6. Wu V, Yeerna H, Nohata N, Chiou J, Harismendy O, Raimondi F, et al. Illuminating the Onco-GPCRome: Novel G protein-coupled receptor-driven oncocrine networks and targets for cancer immunotherapy. The Journal of biological chemistry. 2019;294(29):11062–86.

7. Van Raamsdonk CD, Bezrookove V, Green G, Bauer J, Gaugler L, O’Brien JM, et al. Frequent somatic mutations of GNAQ in uveal melanoma and blue naevi. Nature. 2009;457(7229):599–602.

8. Van Raamsdonk CD, Griewank KG, Crosby MB, Garrido MC, Vemula S, Wiesner T, et al. Mutations in GNA11 in uveal melanoma. N Engl J Med. 2010;363(23):2191–9.

9. Feng X, Degese MS, Iglesias-Bartolome R, Vaque JP, Molinolo AA, Rodrigues M, et al. Hippo-independent activation of YAP by the GNAQ uveal melanoma oncogene through a trio-regulated rho GTPase signaling circuitry. Cancer cell. 2014;25(6):831–45.

10. Ideno N, Yamaguchi H, Ghosh B, Gupta S, Okumura T, Steffen DJ, et al. GNAS(R201C) Induces Pancreatic Cystic Neoplasms in Mice That Express Activated KRAS by Inhibiting YAP1 Signaling. Gastroenterology. 2018;155(5):1593–607 e12.

11. Wilson CH, McIntyre RE, Arends MJ, and Adams DJ. The activating mutation R201C in GNAS promotes intestinal tumourigenesis in Apc(Min/+) mice through activation of Wnt and ERK1/2 MAPK pathways. Oncogene. 2010;29(32):4567–75.

12. Patra KC, Kato Y, Mizukami Y, Widholz S, Boukhali M, Revenco I, et al. Mutant GNAS drives pancreatic tumourigenesis by inducing PKA-mediated SIK suppression and reprogramming lipid metabolism. Nat Cell Biol. 2018;20(7):811–22.

13. Lyons J, Landis CA, Harsh G, Vallar L, Grunewald K, Feichtinger H, et al. Two G protein oncogenes in human endocrine tumors. Science. 1990;249(4969):655–9.

14. Kan Z, Jaiswal BS, Stinson J, Janakiraman V, Bhatt D, Stern HM, et al. Diverse somatic mutation patterns and pathway alterations in human cancers. Nature. 2010;466(7308):869–73.

15. Garcia-Marcos M, Ghosh P, and Farquhar MG. Molecular basis of a novel oncogenic mutation in GNAO1. Oncogene. 2011;30(23):2691–6.

16. Song L, Yu B, Yang Y, Liang J, Zhang Y, Ding L, et al. Identification of functional cooperative mutations of GNAO1 in human acute lymphoblastic leukemia. Blood. 2021;137(9):1181–91.

17. Lyu C, Ye Y, Lensing MM, Wagner KU, Weigel RJ, and Chen S. Targeting Gi/o protein-coupled receptor signaling blocks HER2-induced breast cancer development and enhances HER2-targeted therapy. JCI Insight. 2021;6(18).

18. Lyu C, Ye Y, Weigel RJ, and Chen S. Blocking Gi/o-Coupled Signaling Eradicates Cancer Stem Cells and Sensitizes Breast Tumors to HER2-Targeted Therapies to Inhibit Tumor Relapse. Cancers (Basel). 2022;14(7).

19. Kirui JK, Xie Y, Wolff DW, Jiang H, Abel PW, and Tu Y. Gbetagamma signaling promotes breast cancer cell migration and invasion. J Pharmacol Exp Ther. 2010;333(2):393–403.

20. Tang X, Sun Z, Runne C, Madsen J, Domann F, Henry M, et al. A critical role of Gbetagamma in tumorigenesis and metastasis of breast cancer. The Journal of biological chemistry. 2011;286(15):13244–54.

21. Shajahan AN, Tiruppathi C, Smrcka AV, Malik AB, and Minshall RD. Gbetagamma activation of Src induces caveolae-mediated endocytosis in endothelial cells. The Journal of biological chemistry. 2004;279(46):48055–62.

22. Ma YC, Huang J, Ali S, Lowry W, and Huang XY. Src tyrosine kinase is a novel direct effector of G proteins. Cell. 2000;102(5):635–46.

23. Dbouk HA, and Backer JM. A beta version of life: p110beta takes center stage. Oncotarget. 2010;1(8):729–33.

24. Khalil BD, Hsueh C, Cao Y, Abi Saab WF, Wang Y, Condeelis JS, et al. GPCR Signaling Mediates Tumor Metastasis via PI3Kbeta. Cancer Res. 2016;76(10):2944–53.

25. Ye Y, Tang X, Sun Z, and Chen S. Upregulated WDR26 serves as a scaffold to coordinate PI3K/ AKT pathway-driven breast cancer cell growth, migration, and invasion. Oncotarget. 2016.

26. Ram PT, Horvath CM, and Iyengar R. Stat3-mediated transformation of NIH-3T3 cells by the constitutively active Q205L Galphao protein. Science. 2000;287(5450):142–4.

27. Nautiyal J, Majumder P, Patel BB, Lee FY, and Majumdar AP. Src inhibitor dasatinib inhibits growth of breast cancer cells by modulating EGFR signaling. Cancer Lett. 2009;283(2):143–51.

28. Guarino M. Src signaling in cancer invasion. J Cell Physiol. 2010;223(1):14–26.

29. Martellucci S, Clementi L, Sabetta S, Mattei V, Botta L, and Angelucci A. Src Family Kinases as Therapeutic Targets in Advanced Solid Tumors: What We Have Learned so Far. Cancers (Basel). 2020;12(6).

30. Corre I, Baumann H, and Hermouet S. Regulation by Gi2 proteins of v-fms-induced proliferation and transformation via Src-kinase and STAT3. Oncogene. 1999;18(46):6335–42.

31. Sriram K, Moyung K, Corriden R, Carter H, and Insel PA. GPCRs show widespread differential mRNA expression and frequent mutation and copy number variation in solid tumors. PLoS Biol. 2019;17(11):e3000434.

32. Mayer EL, and Krop IE. Advances in targeting SRC in the treatment of breast cancer and other solid malignancies. Clin Cancer Res. 2010;16(14):3526–32.

33. Myoui A, Nishimura R, Williams PJ, Hiraga T, Tamura D, Michigami T, et al. C-SRC tyrosine kinase activity is associated with tumor colonization in bone and lung in an animal model of human breast cancer metastasis. Cancer Res. 2003;63(16):5028–33.

34. Jallal H, Valentino ML, Chen G, Boschelli F, Ali S, and Rabbani SA. A Src/Abl kinase inhibitor, SKI-606, blocks breast cancer invasion, growth, and metastasis in vitro and in vivo. Cancer Res. 2007;67(4):1580–8.

