## Supplemental Figures for "Active Gαi/o mutants accelerate breast tumor metastasis via the c-Src pathway"

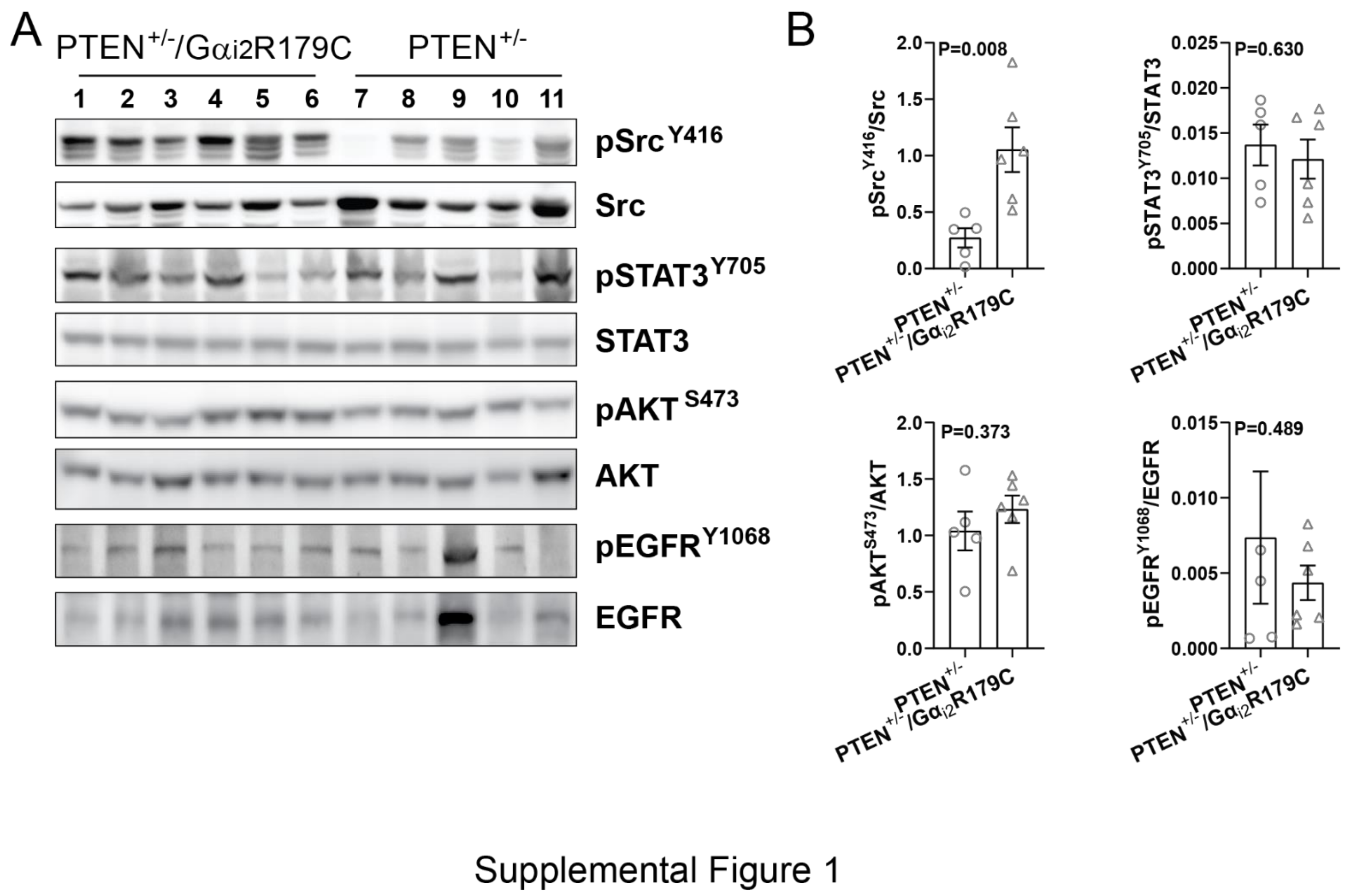


**Supplemental Figure 1. G$\alpha_{i2}$R179C enhances c-Src activation in PTEN$^{+/-}$ tumors. A**, Western blotting showing increased phosphorylation of c-Src$^{Y416}$ and no change in phosphorylation of AKT$^{s473}$, STAT3$^{Y705}$ and EGFR$^{Y1068}$ in PTEN$^{+/-}$/G$\alpha_{i2}$R179C tumors as compared to PTEN$^{+/-}$ tumors. Each lane represents a sample from an individual tumor. **B,** The Western blotting data from **A** were quantified and expressed as the ratio of the phosphorylated to total proteins (**B**). Two tail unpaired Student's *t* test was used for statistical analysis of the data in **B**, and p values are shown.

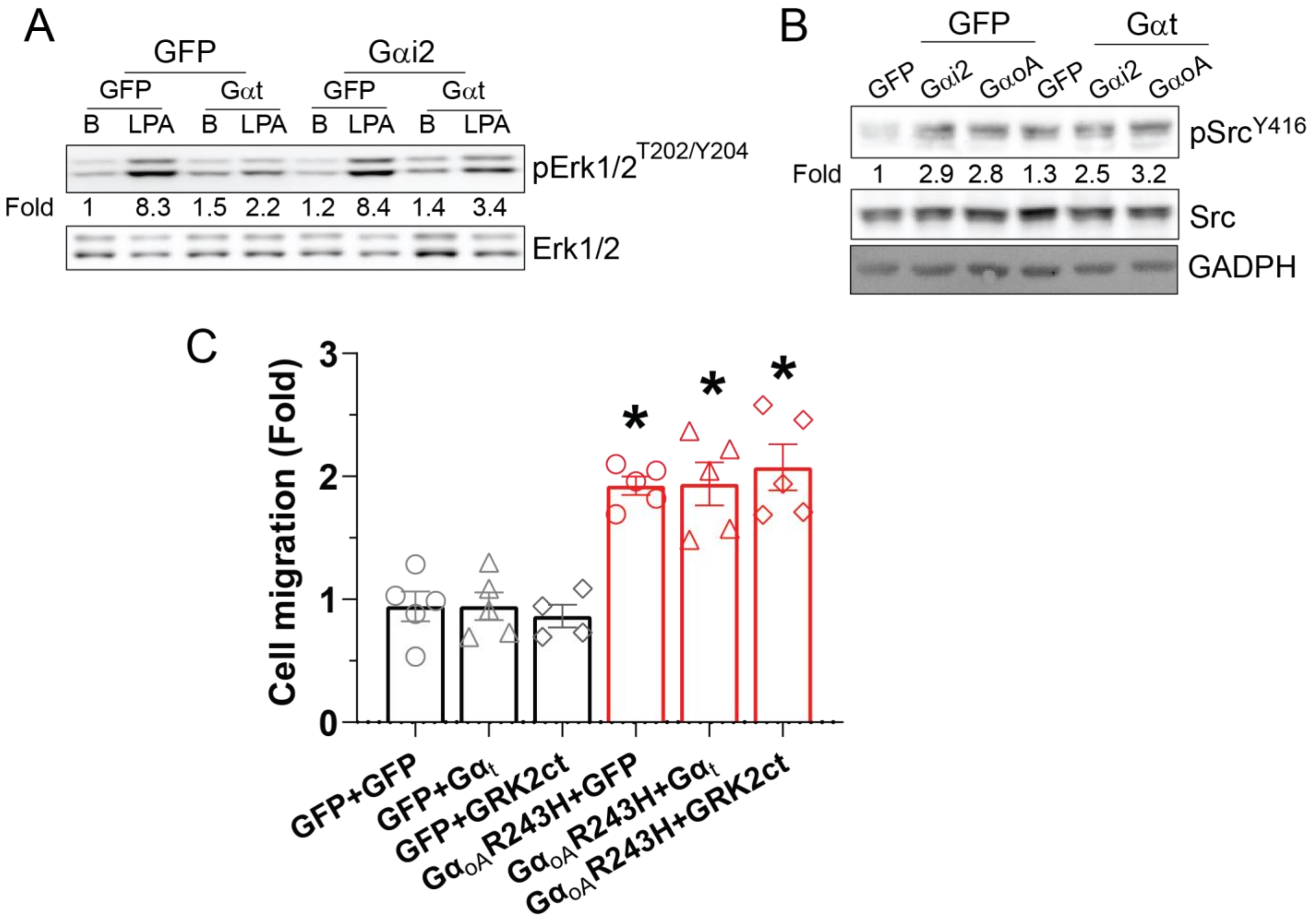


Supplemental Figure 2

**Supplemental Figure 2. Effects of Gβγ scavengers on Gα$_{i2}$R179C- and Gα$_{oA}$R243H-promoted c-Src activation and Neu cell migration. A-B**, Western blotting showing effects of expressing GFP and Gα$_t$ by adenovirus transduction on LPA-stimulated ERK phosphorylation in Neu cells expressing GFP or Gα$_{i2}$R179C (Gα$_{i2}$) (**A**) and c-Src activation in Neu cells expressing GFP, Gα$_{i2}$R179C (Gα$_{i2}$) or Gα$_{oA}$R243H (Gα$_{oA}$) (**B**). **C,** the effect of expressing GFP, Gα$_t$ and GRK2ct by adenovirus transduction on transwell migration of Neu cells expressing GFP or Gα$_{oA}$R243H. *$p<0.05$ *vs* GFP. One-way ANOVA was used for statistical analysis in this Figure.
